# Adapting Lateral Stepping Control to Walk on Winding Paths

**DOI:** 10.1101/2024.07.11.603068

**Authors:** Anna C. Render, Joseph P. Cusumano, Jonathan B. Dingwell

## Abstract

Most often, gait biomechanics is studied during straight-ahead walking. However, real-life walking imposes various lateral maneuvers people must navigate. Such maneuvers challenge people’s lateral balance and can induce falls. Determining how people regulate their stepping movements during such complex walking tasks is therefore essential. Here, 24 adults (12F/12M; Age 25.8±3.5yrs) walked on wide or narrow virtual paths that were either straight, slowly-winding, or quickly-winding. From each trial, we computed time series of participants’ step widths and their lateral body positions relative to their path. We applied our Goal Equivalent Manifold framework – an analysis of how task-level redundancy impacts motor regulation – to quantify how participants adjusted their step width and lateral position from step to step as they walked on these paths. On the narrower paths, participants walked with narrower steps and less lateral position and step width variability. They did so by correcting step-to-step deviations in lateral position more, while correcting step-to-step deviations in step width less. On the winding paths, participants took both narrower and more variable steps. Interestingly, on slowly-winding paths, participants corrected step-to-step deviations in step width more by correcting step-to-step deviations in lateral position less: i.e., they prioritized maintaining step width over position. Conversely, on quickly-winding paths, participants strongly corrected step-to-step deviations in both step width *and* lateral position: i.e., they prioritized maintaining both approximately equally, consistent with trying to maximize their maneuverability. These findings have important implications for persons with gait impairments who may have elevated fall risk.

## INTRODUCTION

Most of what we know about gait biomechanics comes from studying how people walk in straight lines (Richards et al., 2023). However, real-life walking environments are far more complex (Musselman and Yang, 2007), with natural and built features like uneven terrain (Matthis et al., 2018), and various fixed (Twardzik et al., 2019) or moving (Moussaïd et al., 2011) obstacles. In real life, people rarely walk straight-ahead for more than brief periods (Orendurff et al., 2008; Wang and Adamczyk, 2019). Instead, actual walking trajectories are ubiquitously curved (Bergsma et al., 2021; Korbmacher and Tordeux, 2022; Rudenko et al., 2020). While lab-based studies have researched walking along circular arcs of constant (typically ∼1-5m) radii (Courtine et al., 2006; Ho et al., 2023; Rasmussen et al., 2024), real-world walking follows paths of *varying* curvature (Hicheur et al., 2007; Moussaïd et al., 2011; Pham et al., 2011). Walking on curved paths increases lateral instability (Render et al., 2024) and possibly fall risk (Rasmussen et al., 2024). Likewise, older adults often fall while trying to perform such maneuvers, due to inadequate lateral weight-shifting (Robinovitch et al., 2013).

Walking humans naturally exhibit greater instability in the frontal plane (Bauby and Kuo, 2000; McAndrew et al., 2011). Maintaining lateral balance while walking requires controlling center-of-mass motion. Though several mechanisms contribute to this (Fettrow et al., 2019; Harter et al., 2024), the trajectory the center-of-mass follows is most strongly dictated by foot placement (MacKinnon and Winter, 1993; Townsend, 1985). People adjust mediolateral foot placement from step to step to prevent lateral instabilities (Bruijn and van Dieën, 2018; Kuo, 1999). Indeed, foot placement (stepping) is widely considered *the* primary strategy people use to maintain lateral balance (Bruijn and van Dieën, 2018; Hof, 2008; van Leeuwen et al., 2020). Therefore, understanding how people regulate their steps (from step to step) as they walk along non-straight curved walking paths is a critical prerequisite to understanding how they maintain balance, which we analyzed in a companion paper (Render et al., 2024).

Performing any straight or non-straight walking task starts with people taking steps. This involves processes that govern within-step dynamics to ensure viability (i.e., being able to take steps without falling) (Patil et al., 2022, 2024). While viability is necessary to walk, it is not sufficient. People walk with a destination to reach and/or other tasks to achieve (Arechavaleta et al., 2008): i.e., they walk with some purpose. Thus, people must also coordinate their steps to achieve *goal-directed* walking. Among the simplest goals is to follow some designated path (Dingwell et al., 2023; Pham et al., 2011), like a sidewalk or store aisle, etc. Additionally, people also need to adapt their stepping to changes in walking conditions or task demands that arise as they walk, like making lateral maneuvers (Desmet et al., 2022) or avoiding other persons (Olivier et al., 2013).

We developed a framework, based on Goal Equivalent Manifolds (GEMs) (Cusumano and Cesari, 2006; Cusumano and Dingwell, 2013), to propose and test specific hypotheses about how people regulate their steps to walk (Dingwell and Cusumano, 2019; Dingwell et al., 2010). This framework lets us identify the specific stepping variables people prioritize and how they correct errors with respect to proposed target values of those variables. To regulate lateral stepping during straight-ahead walking, our GEM approach identified that people multi-objectively regulate a combination of lateral body position (*z*_*B*_) to remain on their specified path (Dingwell et al., 2023), and step width (*w*) to maintain lateral balance (Dingwell and Cusumano, 2019). We also identified a direct mathematical tradeoff between *z*_*B*_-regulation and *w*-regulation: one cannot easily prioritize one without deprioritizing the other (Dingwell and Cusumano, 2019). During straight-ahead walking, people strongly prioritize *w*-regulation (Dingwell and Cusumano, 2019; Kazanski et al., 2020, 2023).

To walk in different or changing contexts, people must adapt their lateral stepping regulation. Striking the right balance between regulating *z*_*B*_ and *w* depends on the specific walking conditions/task. When people were given explicit task feedback to regulate either *z*_*B*_ or *w* from step to step, they indeed traded off their prioritization by increasing error correction of the feedback task variable, while decreasing error correction of its complement (Render et al., 2021), as we had theoretically predicted (Dingwell and Cusumano, 2019). People similarly traded off these priorities to perform discrete lateral lane changes (Desmet et al., 2022; Desmet et al., 2024), and while walking on continuously narrowing paths (Kazanski et al., 2023). However, when subjected to random external lateral perturbations, people increased error correction of both *z*_*B*_ and *w* (Dingwell et al., 2021; Kazanski et al., 2020). Importantly, none of those experiments assessed walking tasks that implicitly require continuous modulation of both *z*_*B*_ and *w* regulation.

Here, we tasked young healthy adults to navigate various wide or narrow winding paths. Presenting people with paths of different widths challenged their ability to regulate their step widths (*w*). Making these paths oscillate laterally challenged their ability to regulate their lateral position (*z*_*B*_). Thus, we created walking paths that systematically challenged people to weigh how they prioritize both *w* and *z*_*B*_ simultaneously. By amplifying the inherent competition between maintaining *w* and *z*_*B*_, we expected participants to implicitly adapt their lateral stepping regulation to accommodate both factors while navigating these paths. We predicted that on narrower paths, people would trade off decreased *w*-regulation for increased *z*_*B*_-regulation, similar to what people do on continuously narrowing paths (Kazanski et al., 2023). However, as these paths become sufficiently difficult (narrow and oscillating more quickly), we predicted people would simultaneously more strongly correct errors in both *z*_*B*_ and *w*, similar to when they are laterally perturbed (Kazanski et al., 2020).

## METHODS

### Data Availability

All relevant data underlying the results reported here are available on Dryad ([Dataset] Render et al., 2024).

### Participants

Prior to participation, all participants provided written informed consent, as approved by the Institutional Review Board of Pennsylvania State University. Twenty-four young healthy people (Table 1) participated. Participants were screened to ensure no medications, lower limb injuries, surgeries, musculoskeletal, cardiovascular, neurological, or visual conditions affected their gait.

**Table 1.** Participant physical characteristics and assessment scores. Dynamic balance ability and coordination was assessed using the Four-Square Step Test (FSST). Attention and processing speed was assessed using the Four-Choice Reaction Time (4CRT) test. All values except Sex are given as Mean ± Standard Deviation.

| Characteristic: | Value: |
| --- | --- |
| Sex [F/M] | 12 / 12 |
| Age [yrs] | 25.77 ± 3.53 |
| Body Height [m] | 1.74 ± 0.11 |
| Body Mass [kg] | 70.15 ± 14.75 |
| Body Mass Index [kg/m<sup>2</sup>] | 22.97 ± 3.10 |
| Leg Length [m] | 0.93 ± 0.05 |
| **Assessment:** |  |
| FSST [s] | 6.10 ± 1.78 |
| 4CRT [ms] | 384.6 ± 77.9 |

### Assessments

We evaluated participants’ balance using the Four Square Step Test (FSST; Table 1) (Dite and Temple, 2002). Each participant completed three trials and we recorded their mean time. We assessed participants’ attention and processing speed using the Four-Choice Response Task (4CRT; Table 1), administered using PEBL software (https://pebl.sourceforge.net/) (Mueller and Piper, 2014). Each participant completed two blocks of 50 stimuli and we recorded their mean reaction time.

### Experimental Protocol

Participants walked in a M-Gait virtual reality system (Motek Medical, Netherlands; Fig.1A) equipped with a 1.2m wide treadmill. Each participant wore a safety harness secured to the ceiling overhead. For all walking trials, treadmill speed was set to 1.2m/s with matched optic flow speed. Participants initially walked 4-minutes to acclimate to the system.

We tasked participants to walk on paths projected onto the treadmill belt (Fig. 1B). We created pseudo-randomly oscillating paths using a sum of three sin waves with incommensurate frequencies:

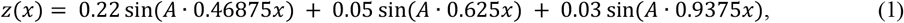

where *z* is the lateral position (in meters) of the path center, A is a frequency scaling factor, and *x* is forward treadmill distance (in meters). We presented participants with each of three oscillation frequency combinations: straight (STR: *A* = 0), “low” frequency (LOF: *A* = 1), and “high” frequency (HIF: *A* = 4) (see Fig. 1A and Supplemental Video). The radii-of-curvature (***R***) of these winding paths varied continuously. LOF paths had a median ***R*** = 32.2m and varied over a 5^th^-to-95^th^ percentile range of 12.6m ≤ ***R*** ≤ 314.6m (i.e., 90% of the total path fell within this range). HIF paths had a median ***R*** = 2.37m and varied over a 5^th^-to-95^th^ percentile range of 0.84m ≤ ***R*** ≤ 23.57m.

**Figure 1.**
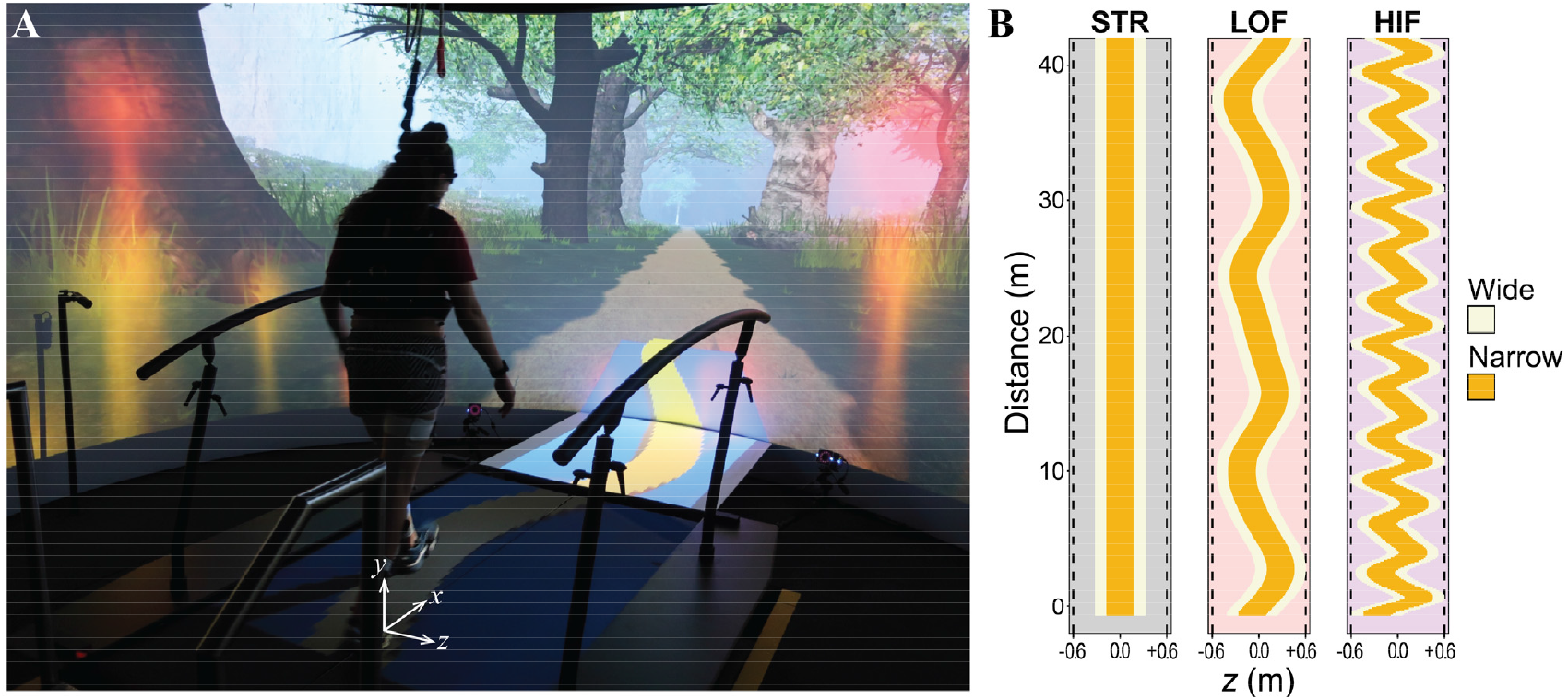
**A)** Photo of a participant walking in the Motek M-Gait virtual reality system, showing a path (here, high-frequency narrow; HIFN) projected onto the treadmill surface. Global coordinates are +*x* forward, +*y* vertical up, and +*z* to the right. See Supplemental Video for examples of participants walking along all 6 paths. **B)** Participants traversed paths of each of three oscillation frequency combinations, Eq. (1): straight (STR; grey panel), low (LOF; red panel), and high (HIF; purple panel). Each path frequency was tested at two different widths, “Wide” (W; path width = 0.6 m; beige) and “Narrow” (N; path width = 0.3 m; yellow). Note the horizontal and vertical axes are not to the same scale (e.g., Narrow HIF looked as shown in **A** when projected). Paths were designed to impose continuously-varying *curved* walking.

Each such path was presented at each of two widths, wide (W = 0.60 m) and narrow (N = 0.30 m), resulting in six unique paths (Fig. 1B). For each, we instructed participants to “stay on the path” and encouraged them to minimize stepping errors. They received visual (firework) and auditory (flute) penalties when steps landed outside the path boundaries (see Supplemental Video). For each condition, participants completed a 1-minute introductory trial, followed by two 4-minute experimental trials. Order of presentation between conditions was counterbalanced across participants using a Latin Square design. To minimize fatigue, participants were allowed to rest as needed after each trial.

### Data Processing

Each participant wore 16 retroreflective markers: four around the head, four around the pelvis (left/ right PSIS and ASIS), and four on each of the left and right shoes (aligned with first and fifth metatarsal heads, lateral malleolus, and calcaneus). Kinematic data were collected at 100 Hz from a 10-camera Vicon motion capture system (Oxford Metrics, Oxford, UK) and post-processed using Vicon Nexus software. Marker trajectories and path data from D-Flow software (Motek Medical, Netherlands) were analyzed in MATLAB (MathWorks, Natick, MA).

We only analyzed the two 4-minute experimental trials for each condition. Marker trajectory data were first low-pass filtered (4th-order Butterworth, cutoff: 10 Hz), then interpolated to 600 Hz to ensure accurate stepping-event detection (Bohnsack-McLagan et al., 2016). Heel strikes were determined using a marker-based algorithm (Zeni et al., 2008).

For each trial, we used the lateral locations of the left and right heel markers to define time series of left and right lateral foot placements. We calculated relevant lateral stepping variables in path-based coordinates (Dingwell et al., 2023) (Fig. 2A). At each step, we aligned the path-based local coordinate system, [*x′, z′*], to be tangent to the point on the centerline of the path closest to the midpoint between the two heel markers for that step. Then, we then defined lateral body position relative to the path center as *z*_*Bn*_*=* ½(*z′*_*Ln*_ *+ z′*_*Rn*_), and step width as *w*_*n*_*= z′*_*Rn*_ – *z′*_*Ln*_ (Fig 2A). This yielded time series of path-aligned {*z*_*Bn*_, *w*_*n*_} (Desmet et al., 2022) for all steps *n* ∈ {1, …, *N*} (Fig. 2B) throughout a trial. For consistency, we analyzed the first *N* = 350 steps of each trial.

**Figure 2.**
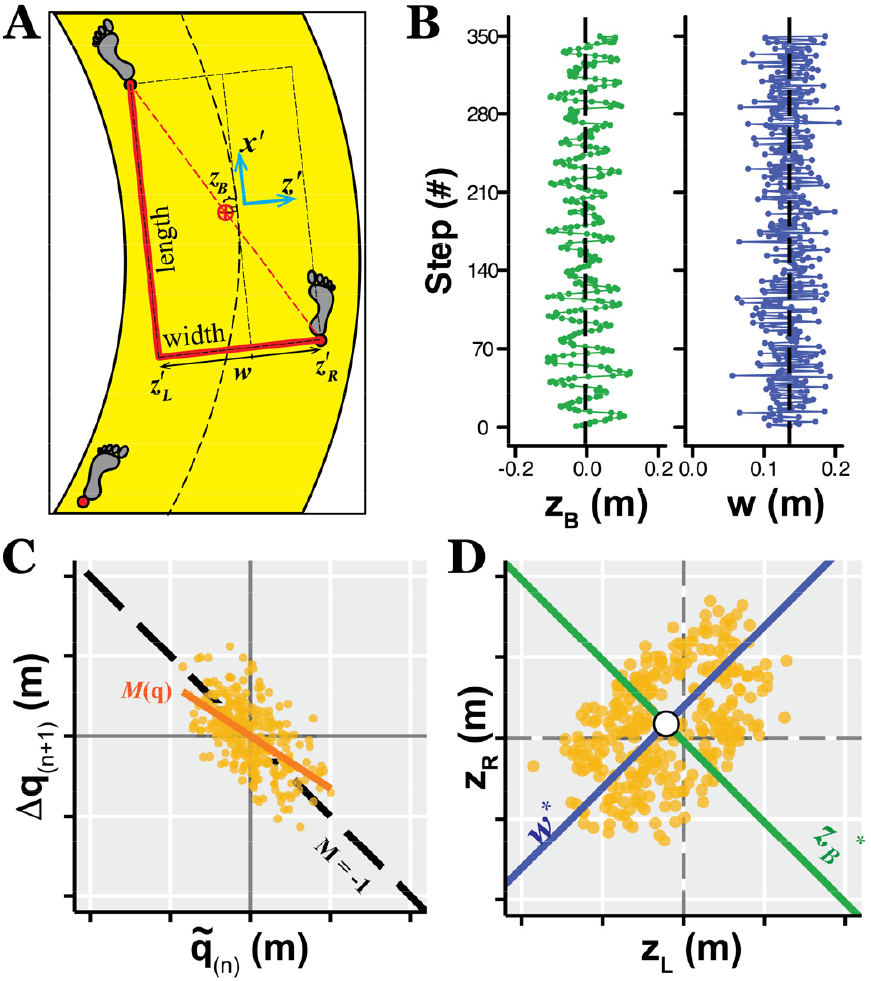
**A**) At each step, we rotate the lab-defined global coordinate system, [*x,z*] to define a path-aligned local coordinate system, [*x′,z′*] in which the *x′* axis is tangent to the centerline of the path closest to the midpoint between the two feet (Dingwell et al. 2023). The locally lateral positions of the left and right feet {*z′*_*L*_, *z′*_*R*_} are used to derive the lateral body position as the midpoint, *z*_*B*_ = ½(*z′*_*L*_ + *z′*_*R*_), and step width as the difference, *w* = *z′*_*R*_−*z′*_*L*_. **B**) Example time series data of path-aligned *z*_*B*_ (green) and *w* (blue) for a representative trial from a typical participant. For the *z*_*B*_ plot (left), the black vertical dashed line indicates the center of the path (i.e., *z*_*B*_ = 0). For the *w* plot (right), the black vertical dashed line indicates the average step width. **C)** Example direct control plot used to quantify how people correct errors in *q* ∈ {*z*_*B*_,*w*} from step-to-step: deviations from the mean on a given step, on, 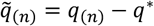, are plotted (yellow markers) against the corrections made on the subsequent step, Δ*q*_(*n*+1)_ *= q*_(*n*+1)_ − *q*_*n*_. The solid yellow diagonal line indicates the error correction slope, *M*(*q*), computed via linear least-squares. The black dashed line indicates “perfect” error correction (*M*(*q*) = −1). **D**) Lateral foot placements (yellow markers) plotted in [*z*_*L*_, *z*_*R*_] plane for a typical trial. Stepping regulation variables [*z*_*B*_, *w*] map onto [*z*_*L*_, *z*_*R*_] as mutually orthogonal GEMs representing constant lateral body position (*z* *= constant; green) and constant step width (*w*\*= constant; blue): position variability, σ(*z*_*B*_) quantifies deviations along *w*^*^, whereas step width variability, σ(*w*), quantifies deviations along *z*_*B*_^*^.

### Stepping Parameters

For each trial performed by each participant under each condition, we first computed within-trial means (*μ*) and standard deviations (*σ*) from the 350-step time series of both lateral body position (*z*_*B*_) and step width (*w*).

We applied two-factor (Frequency × Width) mixed-effects analyses of variance (ANOVA) with repeated measures to test for differences between conditions for each variable: *μ*(*z*_*B*_), *μ*(*w*), *σ*(*z*_*B*_) and *σ*(*w*). We first log-transformed *σ*(*z*_*B*_) and *σ*(*w*) to satisfy normality assumptions. When we found main effects or interaction effects to be significant, we performed Tukey’s pairwise post-hoc comparisons to test for differences between path frequencies (STR, LOF, HIF) for each path width (W, N) and between path widths for each path frequency. We conducted all statistical analyses in Minitab (Minitab, Inc., State College, PA).

### Linear Error Correction

We performed linear error correction analyses (Dingwell and Cusumano, 2015) to quantify the extent to which participants corrected step-to-step deviations in *z*_*B*_ and *w* on subsequent steps. Since we defined *z*_*B*_ and *w* relative to the path (Fig. 2A), they already account for the curving of these paths. We presume people try to maintain (relative to their path) some desired *q*^*^ for each *q* ∈{*z*_*B*_, *w*}. On any given step, *n*, they will exhibit some small deviation from *q*^*^, 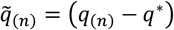. How much they correct those deviations on the subsequent step is then Δ*q*_(*n*+1)_ = *q*_(*n*+1)_ – *q*_(*n*)_. We directly quantified the extent to which 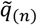 deviations were corrected by Δ*q*_(*n*+1)_ by plotting the corrections (Δ*q*_(*n*+1)_) against their corresponding deviations 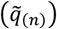 and computing linear slopes, *M(q*), from these plots (Fig. 2C). Slopes equal to −1 indicate deviations that are completely corrected at the next step. Conversely, slopes approaching 0 indicate deviations that go uncorrected (Dingwell and Cusumano, 2015).

We applied the same two-factor (Frequency × Width) mixed-effects analyses of variance (ANOVA) as described above for each variable: *M*(*z*_*B*_) and *M*(*w*). When main effects or interaction effects were found to be significant, we again performed relevant Tukey’s pairwise post-hoc comparisons as described above.

### GEM-Relevant Variance

Any step-to-step regulation of *z*_*B*_ and/or *w* must be enacted by taking steps: *z*_*L*_ and *z*_*R*_. By their definitions (Dingwell and Cusumano, 2019), {*z*_*B*_, *w*} are related to {*z*_*L*_, *z*_*R*_} as:

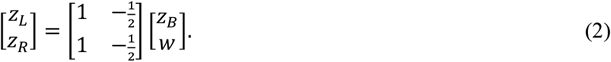

We can thus visualize the effects of the above stepping regulation processes by plotting steps in the [*z*_*L*_, *z*_*R*_] plane (Dingwell and Cusumano, 2019). This [*z*_*L*_, *z*_*R*_] plane exhibits two, mutually perpendicular Goal Equivalent Manifolds (GEMs). One corresponds to maintaining constant position (*z*_B_ *) (Fig. 2D; green), the other to constant step width (*w*\*) (Fig. 2D; blue). We expect how people prioritize their stepping regulation (i.e., error correction) should thus structure the variance in the [*z*_*L*_, *z*_*R*_] plane accordingly. During straight walking, people strongly correct errors in *w* but not in *z*_*B*_, thus exploiting the equifinality available along the *w*\* GEM to exhibit much larger variances in *z*_*B*_ than in *w* (Dingwell and Cusumano, 2019). Conversely, people executing lateral lane change maneuvers exhibit more isotropic variance distributions at their transition step (Desmet et al., 2024). Here, we therefore calculated the ratio *σ*(*z*_*B*_)/*σ*(*w*), where strong prioritization of *w*-regulation over *z*_*B*_-regulation is expected to yield *σ*(*z*_*B*_)/*σ*(*w*) ≫ 1.

We applied the same two-factor (Frequency × Width) mixed-effects analyses of variance (ANOVA) as described above for *σ*(*z*_*B*_)/*σ*(*w*). As warranted, we again performed relevant Tukey’s pairwise post-hoc comparisons as described above.

### Correlations With Baseline Measures

To determine the extent to which baseline tests (FSST and/or 4CRT; Table 1) could predict walking task performance, we correlated these measures against our primary walking outcomes (*M*(*z*_*B*_), *M*(*w*), and *σ*(*z*_*B*_)/*σ*(*w*)) for the Narrow HIF path condition, as this was the most balance-challenging condition presented.

## RESULTS

### Stepping Parameters

Participants exhibited similar mean values of *z*_*B*_ across all conditions (p > 0.18; Fig. 3A; Table 2), thus staying close to the center of each path. On the narrow paths, participants took narrower steps on average, *μ*(*w*), for all path frequency conditions (p < 0.001; Fig. 3B; Table 2). As path oscillation frequency increased (from STR to LOF to HIF) for both path widths, participants exhibited progressively narrower *μ*(*w*) (p < 0.01; Fig. 3B; Table 2).

**Table 2.** Statistical results for differences between path Frequencies (STR, LOF, and HIF) and path Widths (Wide, Narrow) for data shown in Figs. 3-6. ANOVA results (F-statistics and p-values) are provided for main effects of Frequency and Width, and for Frequency × Width interactions. Tukey’s post-hoc comparisons are presented where relevant. Significant differences are indicated in bold. * In cases where Frequency × Width interactions were significant, we considered main effects results to be unreliable and potentially misleading. We drew conclusions instead from the Tukey’s pairwise comparisons, as these account for the effects of the interactions.

| Data In | Variable | Frequency | Width | Frequency × Width | Tukey's (Frequency Effects) | Tukey's (Width Effects) |
| --- | --- | --- | --- | --- | --- | --- |
| Fig. 3A: | $\mu(z_B)$ | $F_{(2,259)} = 1.72$<br>$p = 0.181$ | $F_{(1,259)} = 1.58$<br>$p = 0.210$ | $F_{(2,259)} = 0.60$<br>$p = 0.551$ | N/A | N/A |
| Fig. 3B: | $\mu(w)$ | $F_{(2,259)} = 144.10$<br>$p^* < 0.001$ | $F_{(1,259)} = 411.10$<br>$p^* < 0.001$ | $F_{(2,259)} = 10.24$<br>$p < 0.001$ | STRW-LOFW: $p = 0.003$<br>STRW-HIFW: $p < 0.001$<br>LOFW-HIFW: $p < 0.001$<br>STRN-LOFN: $p = 0.002$<br>STRN-HIFN: $p < 0.001$<br>LOFN-HIFN: $p < 0.001$ | STRW-STRN: $p < 0.001$<br>LOFW-LOFN: $p < 0.001$<br>HIFW-HIFN: $p < 0.001$ |
| Fig. 4A: | $\ln(\sigma(z_B))$ | $F_{(2,259)} = 1944.40$<br>$p^* < 0.001$ | $F_{(1,259)} = 805.84$<br>$p^* < 0.001$ | $F_{(2,259)} = 25.34$<br>$p < 0.001$ | STRW-LOFW: $p < 0.001$<br>STRW-HIFW: $p < 0.001$<br>LOFW-HIFW: $p < 0.001$<br>STRN-LOFN: $p < 0.001$<br>STRN-HIFN: $p < 0.001$<br>LOFN-HIFN: $p < 0.001$ | STRW-STRN: $p < 0.001$<br>LOFW-LOFN: $p < 0.001$<br>HIFW-HIFN: $p < 0.001$ |
| Fig. 4B: | $\ln(\sigma(w))$ | $F_{(2,259)} = 2460.13$<br>$p^* < 0.001$ | $F_{(1,259)} = 233.88$<br>$p^* < 0.001$ | $F_{(2,259)} = 4.87$<br>$p = 0.008$ | STRW-LOFW: $p < 0.001$<br>STRW-HIFW: $p < 0.001$<br>LOFW-HIFW: $p < 0.001$<br>STRN-LOFN: $p < 0.001$<br>STRN-HIFN: $p < 0.001$<br>LOFN-HIFN: $p < 0.001$ | STRW-STRN: $p < 0.001$<br>LOFW-LOFN: $p < 0.001$<br>HIFW-HIFN: $p < 0.001$ |
| Fig. 5B: | $M(z_B)$ | $F_{(2,259)} = 3105.27$<br>$p^* < 0.001$ | $F_{(1,259)} = 107.2$<br>$p^* < 0.001$ | $F_{(2,259)} = 3.67$<br>$p = 0.027$ | STRW-LOFW: $p < 0.001$<br>STRW-HIFW: $p < 0.001$<br>LOFW-HIFW: $p < 0.001$<br>STRN-LOFN: $p < 0.001$<br>STRN-HIFN: $p < 0.001$<br>LOFN-HIFN: $p < 0.001$ | STRW-STRN: $p < 0.001$<br>LOFW-LOFN: $p < 0.001$<br>HIFW-HIFN: $p = 0.002$ |
| Fig. 5D: | $M(w)$ | $F_{(2,259)} = 138.92$<br>$p^* < 0.001$ | $F_{(1,259)} = 128.78$<br>$p^* < 0.001$ | $F_{(2,259)} = 11.13$<br>$p < 0.001$ | STRW-LOFW: $p < 0.001$<br>STRW-HIFW: $p < 0.001$<br>LOFW-HIFW: $p = 0.967$<br>STRN-LOFN: $p < 0.001$<br>STRN-HIFN: $p < 0.001$<br>LOFN-HIFN: $p < 0.001$ | STRW-STRN: $p < 0.001$<br>LOFW-LOFN: $p < 0.001$<br>HIFW-HIFN: $p < 0.001$ |
| Fig. 6B: | $\sigma(z_B)/\sigma(w)$ | $F_{(2,259)} = 272.51$<br>$p^* < 0.001$ | $F_{(1,259)} = 271.93$<br>$p^* < 0.001$ | $F_{(2,259)} = 97.97$<br>$p < 0.001$ | STRW-LOFW: $p < 0.001$<br>STRW-HIFW: $p < 0.001$<br>LOFW-HIFW: $p < 0.001$<br>STRN-LOFN: $p < 0.001$<br>STRN-HIFN: $p = 1.000$<br>LOFN-HIFN: $p < 0.001$ | STRW-STRN: $p < 0.001$<br>LOFW-LOFN: $p < 0.001$<br>HIFW-HIFN: $p = 1.000$ |

**Figure 3.**
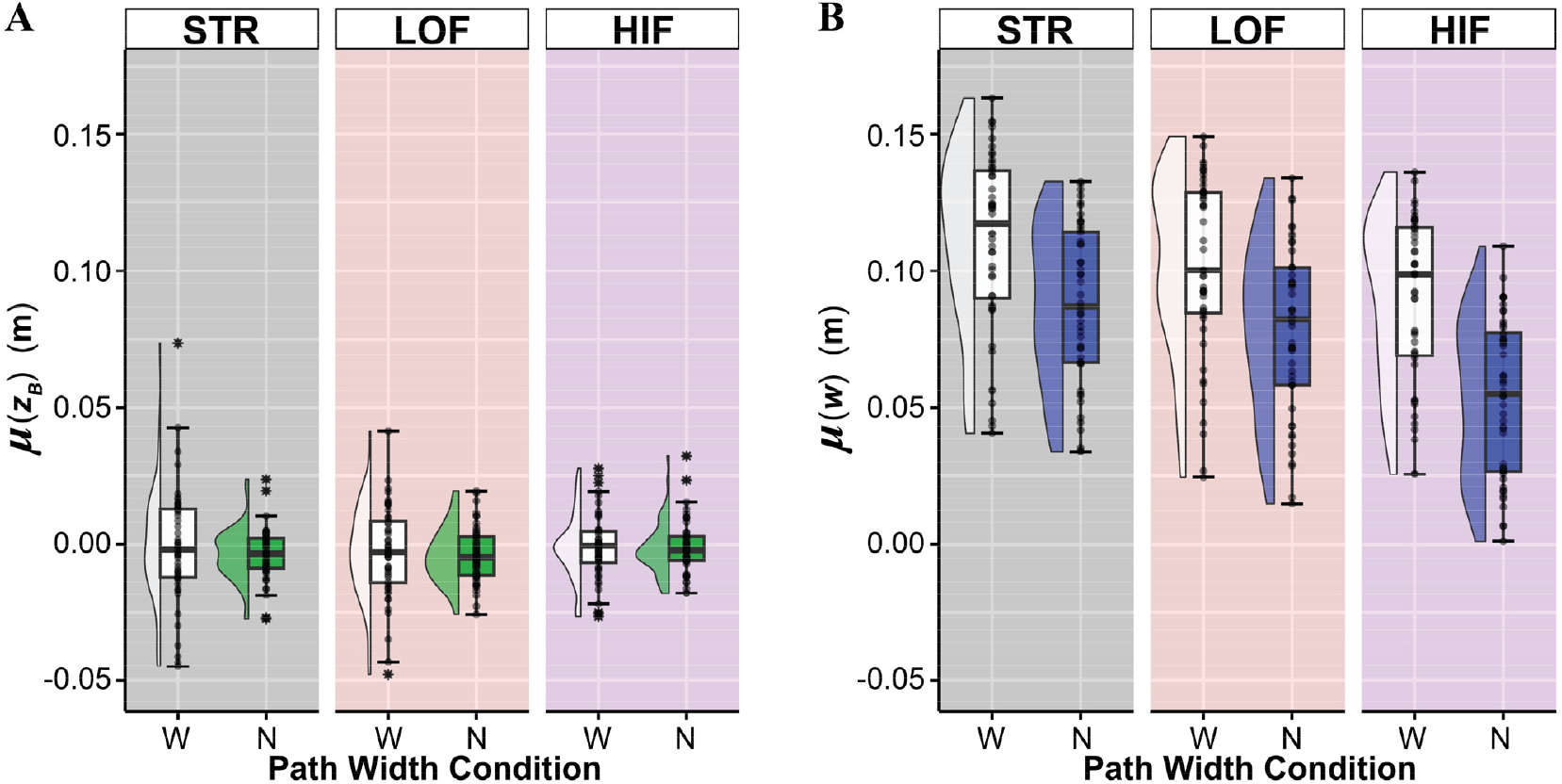
Mean values for (**A**) lateral body position, *μ*(*z*_*B*_), and (**B**) step width, *μ*(*w*), on wide (W) and narrow (N) paths for each path oscillation frequency: STR (grey), LOF (red), and HIF (purple). Box plots show the medians, 1^st^ and 3^rd^ quartiles, and whiskers extending to 1.5 × interquartile range. Values beyond this range are shown as individual asterisks. The overlaid markers are individual data points for each participant from two separate trials. Half-violin plots in each subplot show estimated probability density functions for visual reference. Results of statistical comparisons are in Table 2.

On the narrow paths, participants exhibited significantly less lateral position variability, *σ*(*z*_*B*_) (Fig. 4A), and significantly less step width variability, *σ*(*w*) (Fig. 4B), for all path frequencies (p < 0.001; Table 2). As path oscillation frequency increased, participants exhibited significantly increasing *σ*(*z*_*B*_) (Fig. 4A) and *σ*(*w*) (Fig. 4B), for both path widths (p < 0.001; Table 2).

**Figure 4.**
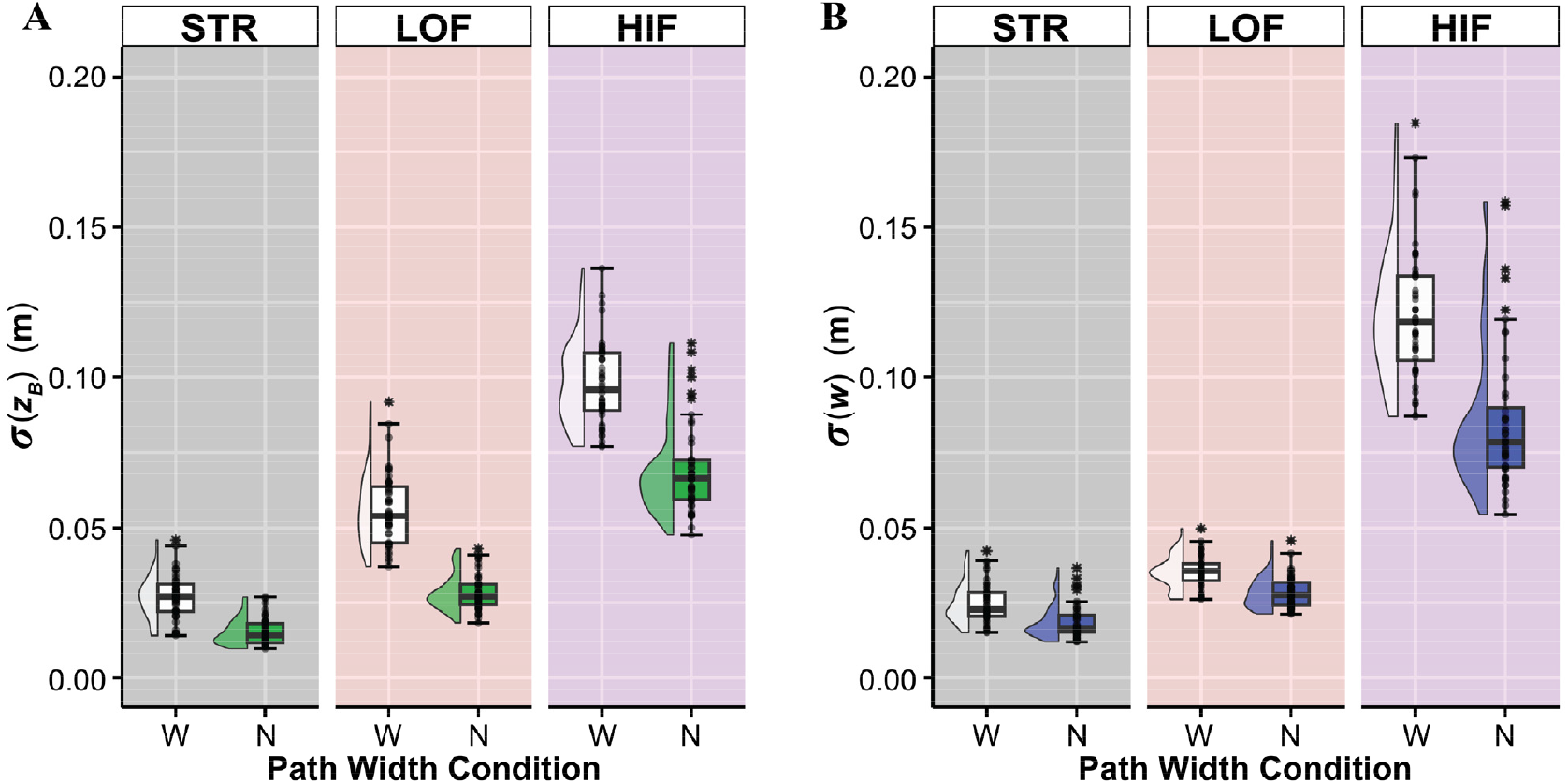
Within-trial standard deviations for (**A**) lateral body position, σ(*z*_*B*_), and (**B**) step width, σ(*w*), on wide (W) and narrow (N) paths for each path oscillation frequency. Data are plotted in the same manner as in Fig. 3. Results of statistical comparisons are in Table 2.

### Linear Error Correction

Across all path frequency and path width combinations, participants consistently exhibited stronger error correction (slopes, *M*, closer to −1.0) for step width (*w*) deviations (Fig. 5C-D) than for lateral position (*z*_*B*_) deviations (Fig. 5A-B), consistent with prior findings (Dingwell and Cusumano, 2019; Kazanski et al., 2023). On the narrow paths, participants corrected position deviations more, *M*(*z*_*B*_) → −1.0, for all path frequencies (p ≤ 0.002; Fig. 5B; Table 2), but corrected step width deviations, *M*(*w*), less so, *M*(*w*) → 0.0, for all path frequencies (p < 0.001; Fig. 5D; Table 2).

**Figure 5.**
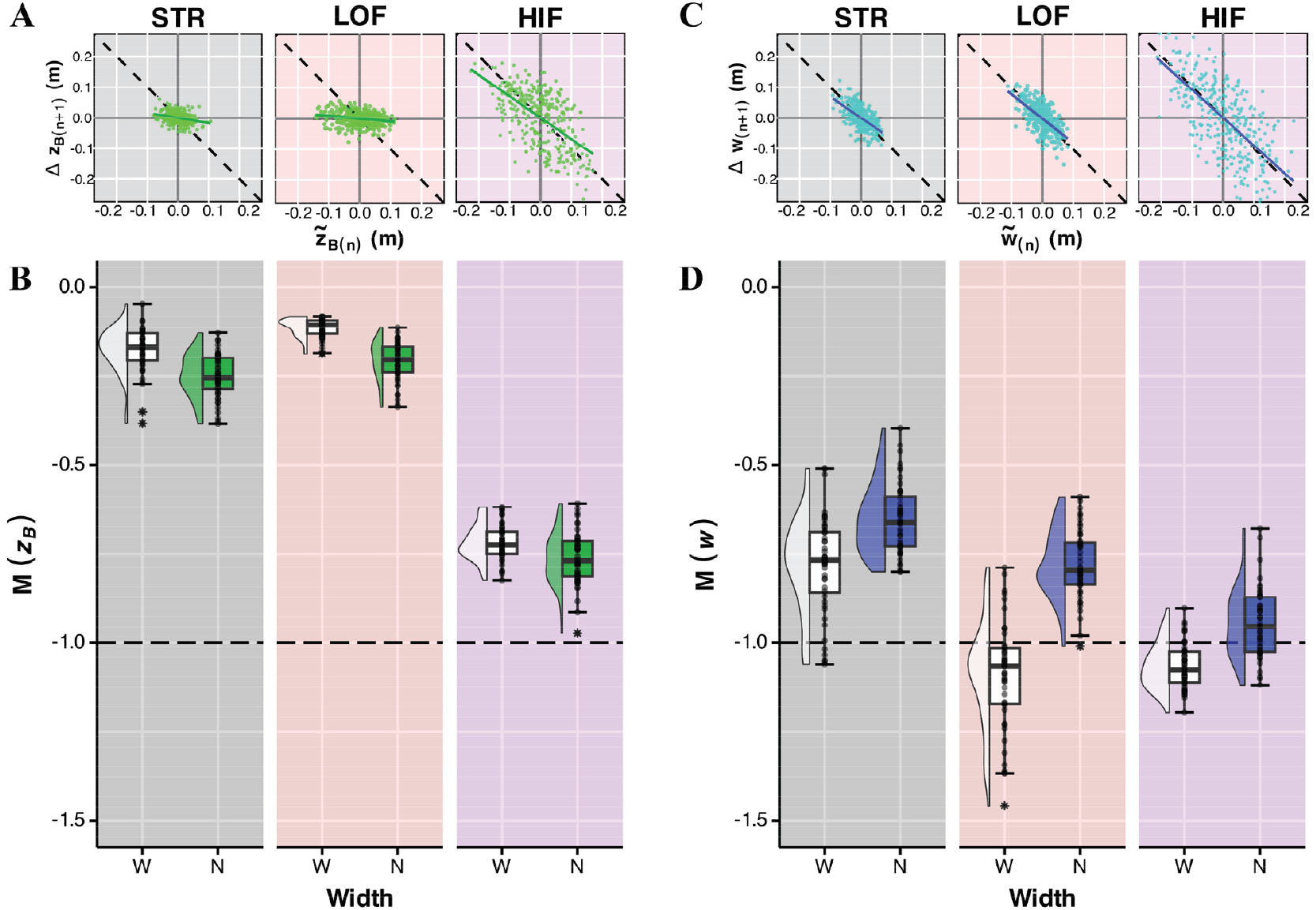
Representative direct control plots of 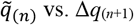 (see Fig. 2C) from a typical participant walking on the three different Wide paths for (**A**) lateral body position (*q* ≡ *z*_*B*_) and (**C**) step width (*q* ≡ *w*). Group data for linear error correction slopes for (**B**) lateral body positions, *M*(*z*_*B*_), and (**D**) step widths, *M*(*w*), on the wide (W) and narrow (N) paths for each oscillation frequency: STR (grey), LOF (red), and HIF (purple) for all trials and participants, plotted in the same manner as in Figs. 3-4. Horizontal dashed lines at slope *M* = −1 indicate perfect error correction: i.e., values above this line reflect *under*-correction, whereas values below this line reflect *over*-correction. Results of statistical comparisons are in Table 2.

As path oscillation frequency increased from STR to LOF paths, participants corrected *z*_*B*_ deviations less (↑ *M*(*z*_*B*_)) for both path widths (p < 0.001; Fig. 5B; Table 2) and corrected *w* deviations more (↓ *M*(*w*)) for both path widths (p < 0.001; Fig. 5D; Table 2). As path oscillation frequency increased further from LOF to HIF paths, participants corrected *z*_*B*_ deviations much more (↓ *M*(*z*_*B*_)) for both path widths (p < 0.001; Fig. 5B; Table 2) and also corrected *w* deviations more (↓ *M*(*w*)) for the narrow path (p < 0.001), but not for the wide path (p = 0.967) (Fig. 5D; Table 2).

### GEM-Relevant Variance

On the wide STR and LOF paths, participants much more strongly corrected step width deviations (Fig. 5D) than position deviations (Fig. 5B). As expected (Dingwell and Cusumano, 2019), this yielded variance distributions in the [*z*_*L*_, *z*_*R*_] plane strongly aligned with the *w*^*^ GEM: i.e., *σ*(*z*_*B*_)/*σ*(*w*) ≫ 1 (Fig. 6).

**Figure 6.**
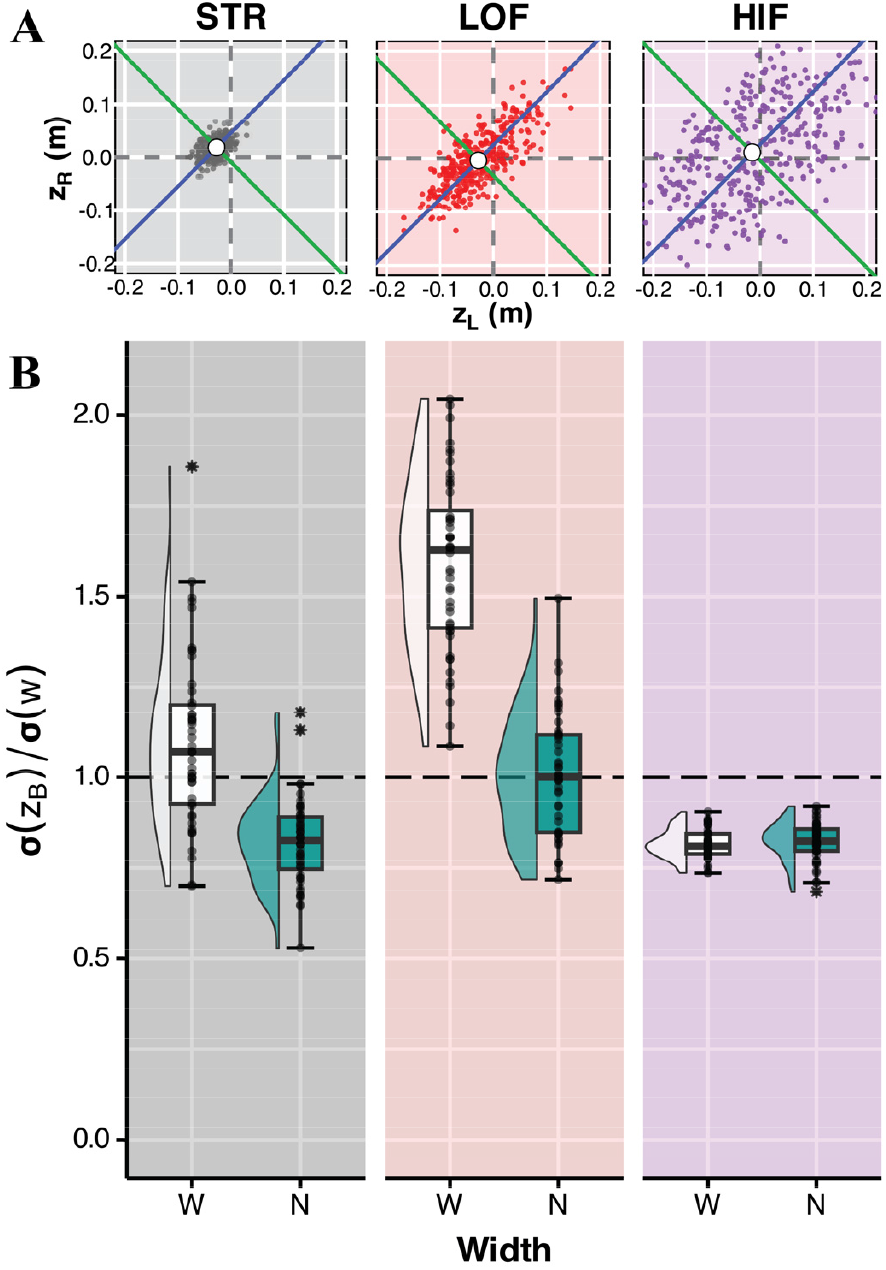
**A**) Example lateral foot placements plotted in [*z*_*L*_, *z*_*R*_] plane (see Fig. 2D) from a typical participant walking on the Wide paths. **B**) Group data for the ratio of lateral body position to step width variances, σ(*z*_*B*_)/σ(*w*), for all trials and participants, plotted in the same manner as in Figs. 3-4. Horizontal dashed lines at σ(*z*_*B*_)/σ(*w*) = +1 indicate isotropic variance distributions. Results of statistical comparisons are in Table 2.

On the narrow paths for these STR and LOF frequency conditions, however, as participants decreased their *w*-regulation to increase *z*_*B*_-regulation (Fig. 5), their variance distributions became significantly more isotropic (p < 0.001; *σ*(*z*_*B*_)/*σ*(*w*) → 1; Fig. 6; Table 2).

On the HIF paths, as participants strongly corrected deviations in both lateral position (Fig. 5B) and step width (Fig. 5D), their variance distributions became significantly more isotropic (*σ*(*z*_*B*_)/*σ*(*w*) < 1) than for the wide STR path (p < 0.001; Fig. 6; Table 2) or either LOF path (p < 0.001; Fig. 6; Table 2).

### Correlations With Baseline Measures

Having faster FSST times predicted, for the Narrow HIF path condition, stronger lateral position error correction (↓ *M*(*z*_*B*_); R^2^ = 31.8%, p = 0.004), coupled with somewhat weaker step width error correction (↑ *M*(*w*); R^2^ = 8.5%, p = 0.2). However, this did not alter the variance distributions (*σ*(*z*_*B*_)/*σ*(*w*); R^2^ = 0.2%, p = 0.8) (Fig. 7). Having slower 4CRT reaction times (not shown) predicted qualitatively similar (though somewhat less statistically significant) performance in this Narrow HIF path walking task.

**Figure 7.**
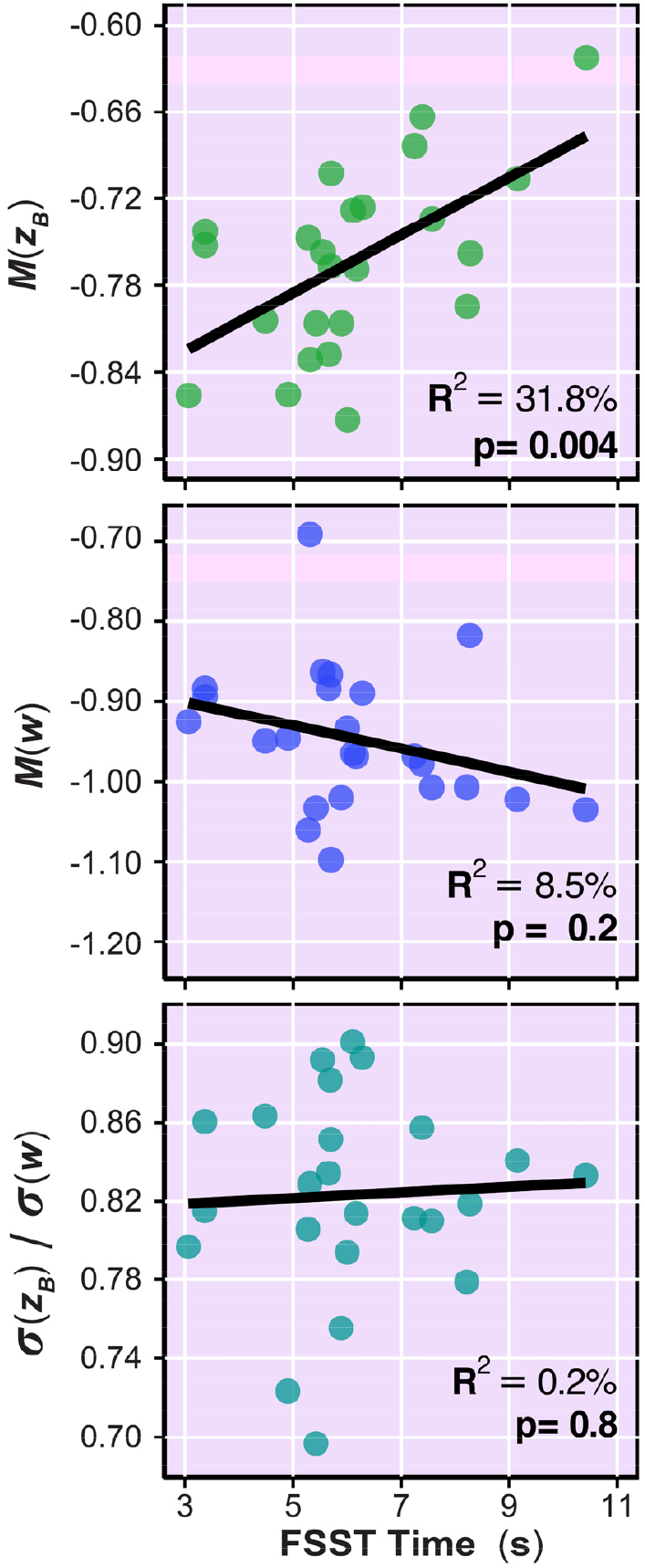
Linear correlations between participants’ baseline average Four-Square Step Test (FSST; Table 1) times and their average *M*(*z*_*B*_) slopes, *M*(*w*) slopes, and σ(*z*_*B*_)/σ(*w*) ratios when walking in the Narrow HIF path condition. In each plot, single markers indicate the average values for each participant. Black solid straight lines indicate linear best fit lines. Correlation strengths (R^2^) and statistical significance (p-values) for each relationship are indicated on each sub-plot.

## DISCUSSION

Being able to walk around curves or through narrow or crowded spaces are important for our daily mobility (Musselman and Yang, 2007). Walking is goal-directed (Arechavaleta et al., 2008) and people readily modify their stepping to navigate specific environmental conditions (Bergsma et al., 2021; Twardzik et al., 2019) and tasks (Acasio et al., 2017; Desmet et al., 2024; Kazanski et al., 2023; Render et al., 2021). Here, we asked people to walk on continuously-winding pathways. We aimed to determine how people trade off the competing task goals, maintain step width or lateral position, in environments designed to demand both simultaneously. We applied our GEM framework for lateral stepping to quantify how people modify stepping regulation and the resulting variance structures on both narrow and winding walking paths.

When walking on the Narrow paths, participants took narrower steps (Fig. 3B) similar to (Schrager et al., 2008), with less lateral position variability (Fig. 4A) and less step width variability (Fig. 4B). Results confirmed our first prediction that, on the narrower paths, people would trade off decreased *w*-regulation for increased *z*_*B*_-regulation, as the GEM framework predicts (Dingwell and Cusumano, 2019), and as we found previously for continuously-narrowing paths (Kazanski et al., 2023). Specifically, people corrected deviations in their lateral position *more* (Fig. 5B), while correcting deviations in step width *less* (Fig. 5D). That people made this trade-off was also reflected in the structure of their stepping variance in the [*z*_*L*_, *z*_*R*_] plane (Fig. 6), where for the STR and LOF conditions particularly, the σ(*z*_*B*_)/σ(*w*) variance ratios decreased closer to 1 on the narrower paths.

Path sinuosity also altered lateral stepping. While navigating more sinuous paths, people took both narrower (Fig. 3B) and more variable (Fig. 4A-B) steps. Interestingly, on the LOF winding paths, people traded-off correcting deviations in their step width *more* (Fig. 5C-D) at the expense of correcting deviations in their lateral position *less* (Fig. 5A-B). This was also reflected in the *σ*(*z*_*B*_)/*σ*(*w*) ratios that were higher in LOF than STR (Fig. 6A-B). Thus, on these moderately-challenging LOF winding paths, people increased their prioritization of maintaining step width, likely to maintain lateral balance (Render et al., 2024).

Consistent with our second prediction, on more sinuous winding paths (HIF), people more strongly corrected errors in both lateral position (Fig. 5A-B) *and* step width (Fig. 5C-D), very similar to what people do when subjected to external (mechanical or visual) lateral perturbations (Kazanski et al., 2020). Critically, however, in (Kazanski et al., 2020), those perturbations were both unpredictable and externally imposed. Conversely, here, people could see their upcoming paths well in advance (Matthis et al., 2017), so their stepping movements were both readily predictable and internally planned (Drew and Marigold, 2015). Our findings thus demonstrate that whether reactive or anticipatory, people adopt similar stepping regulation strategies both when responding to external perturbations and when executing pre-planned tasks.

That people adopted these stepping regulation strategies was also reflected in the variance structure of their stepping data plotted in the [*z*_*L*_, *z*_*R*_] plane (Fig. 6), where on the HIF paths, their σ(*z*_*B*_)/σ(*w*) variance ratios were much closer to 1 for both path widths. Similarly close-to-isotropic [*z*_*L*_, *z*_*R*_] plane variance distributions were observed when people performed discrete lateral maneuvers (Desmet et al., 2024) or walked on narrowing paths (Kazanski et al., 2023). Thus, on these more-challenging HIF winding paths, people prioritized maintaining both step width *and* lateral position to similar degrees, likely to maximize their maneuverability (Acasio et al., 2017; Desmet et al., 2022).

One cannot study control without considering task goals, since “control” signifies the process of correcting errors that deviate from them. Here, we asked people to “stay on the path”. We therefore chose *z*_*B*_* as a *candidate* path goal (Dingwell et al., 2023), and *w*\* as a *candidate* step width goal. Our analyses then test the extent to which step-to-step fluctuations in the data are consistent with this hypothesis. Deviations from *z*_*B*_* and *w*\* were ≤ ∼5-12cm on the Narrow (30cm) paths and ≤ ∼8-15cm on the Wide (60cm) paths (Fig. 4). Hence, participants deviated little from these proposed targets. Likewise, while deviations in *z*_*B*_ were largest for the HIF paths (Fig. 4A), participants also corrected those deviations most strongly (Fig. 5B). Conversely, on the STR and LOF paths, while participants corrected *z*_*B*_ deviations less quickly (Fig. 5B), they also exhibited much smaller deviations initially (Fig. 4A). Together, these findings confirm that *z*_*B*_* and *w*\* are indeed plausible stepping goals for these winding paths, consistent with straight walking (Dingwell and Cusumano, 2019; Dingwell et al., 2010) and other non-straight (Desmet et al., 2022) walking tasks.

However, participants might still have chosen to follow paths somewhat different from those presented. Multiple ways have been proposed to predict a person’s “ideal” path in a given environment (Rudenko et al., 2020), using physics-based models (Helbing et al., 2005), heuristic rules (Moussaïd et al., 2011), or various optimization criteria (Arechavaleta et al., 2008; Brown et al., 2021; Pham et al., 2007). Any of these *could* yield different candidate paths. As we cannot know it for certain, each then becomes a testable hypothesis regarding the person’s true intended path. To our knowledge, no one has attempted to assess how people regulate stepping relative to paths estimated by any of these other approaches. Future work should explore such questions.

The four-square step test (FSST) is a specific functional clinical test of lateral maneuverability (Dite and Temple, 2002). Here, those participants who executed faster FSST times also corrected position (*z*_*B*_) errors more and step width (*w*) errors less (Fig. 7). Hence, those who exhibited stepping patterns that more effectively traded-off stability for maneuverability (Desmet et al., 2022; Desmet et al., 2024) also exhibited faster FSST times. The correlations shown in Fig. 7 thus offer strong evidence of the *concurrent validity* (Portney and Watkins, 2015) of the GEM motor regulation framework and of these lateral stepping regulation measures. Greater ability to adapt one’s lateral stepping regulation to enhance maneuverability appears an important contributor to being able to perform functional tests like the FSST faster. The analyses presented here thus reveal important stepping regulation mechanisms people use to achieve better performance in such functional tasks.

For individuals with gait impairments, many prior studies and clinical training programs have subjected individuals to perturbations (Mansfield et al., 2015). However, based on findings from (Robinovitch et al., 2013), circumstances of falls do not always involve external perturbations. Often, people fall from negotiating a maneuver improperly (Ambrose et al., 2013). Our findings here support that walking paradigms that vary path width and sinuosity may offer more ecologically valid and effective gait interventions to target specific adaptive strategies that improve overall stepping performance and mobility, which may in turn help improve lateral balance (Render et al., 2024).

## ACKNOWLEDGEMENTS

The authors thank Dr. David M. Desmet and Dr. Meghan E. Kazanski for their contributions and technical support throughout data collections. This work was supported by NIH grants 1-R01-AG049735 & 1-R21-AG053470 to JBD & JPC.

## Notes

**CONFLICT OF INTEREST:** The authors declare that there is no conflict of interest associated with this work.

https://doi.org/10.5061/dryad.3tx95x6rb

### Competing Interest Statement

The authors have declared no competing interest.

### Summary of Updates

Updated analyses and results. Revised manuscript in response to reviewer comments.

